# Mimicking and mitigating the cutaneous response to transcranial electrical stimulation using interferential and combinatorial techniques

**DOI:** 10.1101/2021.08.31.456394

**Authors:** Bryan Howell, Cameron C. McIntyre

## Abstract

Transcranial electrical stimulation (tES) is a promising adjunct treatment for neurological impairment and mental health disorders. The modulatory effects of tES are small to moderate, and accrue over days to weeks with repeated administration, but these effects are also inconsistent across individuals, which poses a challenge for its clinical administration. Some of the variability in tES may stem from uncontrolled behavioral factors, and inadequate dosing of current across individuals, so new strategies are needed to address these issues. We evaluated the biophysics of emerging techniques for tES and provided new testable hypotheses for the tolerability of interferentail and combinatorial waveforms. Millisecond pulsatile currents may serve as suitable alternatives to alternating currents in modulating neural spike timing from tES. Pulsatile currents limit spike generation in nerves and may be tolerated above the standard limit of 2 mA when combined with a direct current to block nerve activation. Additionally, we posit that combinations of kilohertz interferential currents can mimic the nerve response of different tES waveforms but with minimal modulation of cortical neurons, providing a new strategy for active placebo stimulation. These results will help guide design of interferential tES strategies for better blinding and provide a testable model for evaluating the tolerability of new combinatorial strategies.

## Introduction

Transcranial electrical stimulation (tES) is an evolving adjunct therapy for movement ^1–3^ and psychiatric ^4–6^ disorders. Conventional tES passes up to 2 mA of electrical current across the cranium, generating weak electric fields of ∼1 V/m in the cortex ^7^. Electric fields of this magnitude are not strong enough to initiate action potentials in neurons, but these fields can modulate spike timing, indirectly, by dynamically altering the neuron’s membrane potential ^8,9^. The safety and tolerability of tES ^10^ make it a promising tool for domiciliary care ^11–13^. However, the prospects of tES as a reliable therapy remain hampered by the substantial variability in behavioral effects within and across individuals ^14–16^.

Insufficient blinding of tES is one major source of variability in outcomes ^17^. Sham tES applies two triangular ramps of current at the beginning and end of sessions ^18^, and for the remaining time, stimulation is either off or at a very small dose (e.g., 0.1 mA) ^19^. Sham tES provides a carryover effect from transient nerve activation ^18,20^, implicating psychological effects from expectations ^21^, but the efficacy of this blind wanes with increasing experience ^22^ and at larger doses ^23^. Moreover, tES evokes reddening of the skin (i.e., erythema), which is difficult to mask ^24^; and tES activates sensory nerves ^25,26^, which can also impact cortical excitability ^27^. Dermatological pretreatment has been proposed to mitigate sensation and erythema ^28,29^, but this comes at the cost of complicating the clinical administration of tES. Alternatively, current steering between multiple electrode pairs has been proposed to minimize electric field intensities in the target brain region ^30–33^. This “active” sham strategy has the advantage of preserving a cutaneous response to tES, but the introduction of multiple foci of stimulation has its own caveats, as modulation of other brain regions and nerves may introduce other sources of latent variability.

Insufficient dosing is another major limiting factor of tES due largely to anatomical and physiological differences across individuals ^34,35^. Thicker layers of CSF and skull ^36,37^, and a more resistive dermis ^38,39^ shunt more current away from the brain, causing field reductions of 100% or more across individuals at fixed doses ^40^. These field fluctuations are further exacerbated by movement or misplacement of electrodes ^41,42^. Therefore, a fixed maximum dose of 2 mA may not achieve electric fields of even 1 V/m in many subjects ^37^. Dosimetric simulations combining current flow modeling and electrophysiological recordings predict that larger currents as large as 3–6 mA are required to achieve field strengths above 1 V/m in most individuals ^43^, but dosing of this magnitude may not be tolerable with standard techniques.

Higher dosing with tES may be achievable with unconventional techniques, such as transcranial random noise stimulation (tRNS) ^44^, transcranial pulsed current stimulation (tPCS) ^45^, and transcranial temporally interfering stimulation (tTIS) ^46^. These alternative techniques vary in time similar to tACS; however, their amplitudes can be modulated, and frequencies typically range from hundreds to thousands of hertz. Because neurons can be approximated as low-pass filters, increasing the stimulus frequency (or reducing the stimulus duration) is expected to diminish its polarizing strength ^47^, but at very high frequencies, other physiological effects may also emerge, which are not fully understood ^48^. Additionally, different techniques can be combined to produce unique modulatory effects different from those of the constituents ^49,50^, but the biophysics of these approaches has not been thoroughly studied.

Our goal in this study was to evaluate the biophysics of emerging techniques for tES both at the scalp and cortical levels. Temporally interfering currents are amplitude-modulated and contain high frequencies similar to tRNS, so our analyses focused on the comparative performance of tPCS and tTIS against standard techniques, including tACS and tDCS. Our results help explain why the cutaneous response to tACS declines considerably above hundreds of hertz ^51,52^, why kilohertz tACS (and by extension, tTIS) is minimally perceivable ^53^, and why tDCS is tolerable at relatively large doses above the standard limit of 2 mA ^54,55^. From these insights, we propose two testable novel strategies. First, a combination of tPCS and tDCS to coordinate spike timing but with large fibers blocked due to inactivation of their sodium channels. Second, an active placebo control that uses tTIS to activate cutaneous nerves but with negligible polarization of cortical neurons. Together, these results highlight the potential for future model-driven design of tES based on established biophysical principles.

## Materials and methods

### Modeling tES of nerves

Current-regulated tES of the scalp was modeled with an ideal disk electrode in contact with a bulk tissue medium. The disk electrode was 3 inches (76.2 mm) in diameter with a return electrode at infinity, so the electric potentials were approximated using the following analytic expression ^56–58^:

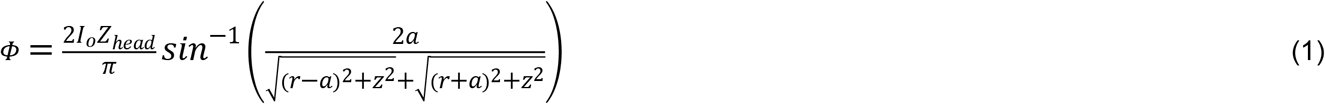

where *Φ* is the potential field, *I*_*o*_ is the applied current, *Z*_*head*_ is the equivalent bulk tissue impedance of the head, *a* is the radius of the disk electrode, and *r* is the radial distance from the disk center.

We estimated *Z*_*head*_ using a seven-domain head model consisting of six concentric spherical shells (Domains 1–6) surrounding a spherical brain (Domain 7). The geometry of the Domains was consistent with the geometry of an average adult human head. Domain 1 combined dermis and epidermis (with a layer thickness of 1.2 mm) ^59^, Domain 2 combined subdermis and fat (1 mm), Domain 3 was muscle (2 mm), and Domain 4 was tendon (0.6–1.5 mm) ^60^. Domain 5 was cranium (7.7 mm) ^61^, Domain 6 was cerebral spinal fluid (CSF, 2.5 mm) ^62^, and Domain 7 was the brain with a radius of ∼74 mm. Additional circular domes (9.5 mm in thickness and ∼76.2 mm in arc length) were placed against Domain 1 at the nadir and zenith to represent saline-soaked sponges, which we refer to as Domains E1 and E2, respectively. Electrodes were the external boundaries of Domains E1 and E2.

Electrical properties were defined using parametric models of dielectric (Cole-Cole) dispersion ^63^,

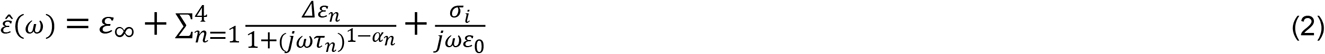

where *Δ*ε = ε_∞_– ε_s_, ε_∞_is the permittivity at large frequencies (*ωτ* >> 1), ε_s_is the permittivity at small frequencies (*ωτ* << 1), ε_0_ is the permittivity of free space, *j*^2^ = –1, *ω* is the angular frequency, *τ* is the dispersion time constant, *α* controls the broadening of each dispersion, *σ*_*i*_is the static ionic conductivity at *ω* = 0, and *n* is the dispersion number. The four dispersions were alpha (*n*=1), beta (*n*=2), low gamma (*n*=3), and medium gamma (*n*=4). Equation 2 was re-expressed in terms of an equivalent permittivity (ε) and conductivity (*σ*),

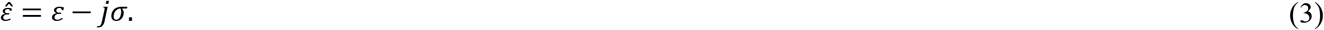

ε and *σ* were applied to each respective domain, for each *ω*. Therefore, ε was the real part of Equation 2 multiplied by ε_0_; and *σ* was the negative imaginary part of Equation 2 multiplied by ε_0_and *ω*. Each domain was assigned one set of material properties using the measurements taken by Gabriel and Gabriel ^63^. Domain 1 was skin (wet); Domain 2 was fat (infiltrated); Domain 3 was muscle; Domain 4 was tendon; Domain 5 was bone (cortical); Domains 6, E1, and E2 were CSF; and Domain 7 was brain (grey matter). CSF was modeled as purely conductive (1.5 S/m, ^64^), whereas all other tissue domains were modeled as “lossy” mediums with frequency-dependent electrical properties. The parameters of Equation 2 for each respective tissue type are found in the work by Gabriel and Grabiel ^63^. All external boundaries were treated as ideal insulators with zero current flux except those representing the two electrodes.

The head impedance (*Z*_*head*_) was calculated numerically. First, 1 mA of current was passed between two, three-inch diameter disk electrodes at the zenith and nadir of the head model. Second, we solved Laplace’s equation for the electric potentials in the tissue medium,

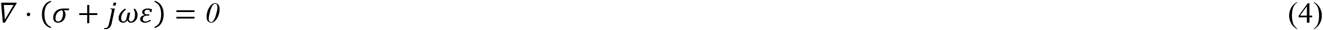

Where, *σ* and *ε* varied by frequency and domain. *Z*_*head*_was calculated using Ohm’s law (V = IR). Non-sinusoidal voltage waveforms were calculated using a Fourier approach ^65^ with 500,000 frequencies uniformly sampled between 0 and 100 kHz. We calculated the complex Fourier transform coefficients at each frequency, scaled each coefficient by its respective impedance, and used the inverse Fourier transform to obtain the applied voltage waveform, and then *Z*_*head*_. We ignored the additional filtering effects of the electrode-tissue interface.

A model nerve 2 mm in diameter was constructed for simulating electrocutaneous stimulation. The nerve’s axis was oriented in the *x* direction and placed 7 mm below the scalp’s surface at the zenith. Two sets of 300 axons were distributed uniformly and randomly within the cylindrical region. For each set of axons, two uniform distributions of radii (between 0 and 1) and angles (between 0 and 2*π*) were generated using latin hypercube sampling to ensure a near-random and uniform distribution of values across each respective range. The square roots of the radii were used when transforming cylindrical coordinates to Cartesian coordinates, to preserve the uniform spacing of the points within the nerve’s cross-section. Axon were seeded at these points in the yz plane transverse to its axis. 150 fiber diameters were randomly assigned to five integer values between 8 μm and 12 μm with proportions of 8:37:60:37:8 from smallest to largest, and this was repeated with another independent random sample but for five fiber diameters between 2 μm and 6 μm. The population of larger fiber diameters represented A*β* cutaneous afferents, whereas the smaller fibers represented A*δ* nociceptive afferents (Fig. 5A).

Axonal responses to extracellular stimulation were simulated using a model of a mammalian motor axon in the Neuron (v7.3) simulation environment ^66^. The geometrical and physiological parameters of the axon model were those originally published by McIntyre et al. ^67^ but with a few notable changes to the geometric parameters. One, geometric parameters not explicitly defined in the original paper were interpolated as a function of fiber diameter using fourth-order polynomials fit to the published data ^68^. Two, we assumed a linear relationship between fiber diameter and internodal lengths for fiber diameters of 4 *μ*m or less ^69^. And three, for these same small fibers, paranodal lengths were 4 % of the internodal length ^70^. Simulations were conducted using Backward-Euler with a uniform time step of 2.5 *μ*s except for cases where stimulation carrier frequencies were greater than 5 kHz (or pulse widths were shorter than 0.1 ms), in which case, a step size of 1 *μ*s was used. Simulations were run for at least 40 ms, and propagation of action potentials was measured at the axons’ termini. Axons were quiescent if they initiated no action potentials or responded only transiently. Axons were active if they fired one or more action potentials for each depolarizing phase of the current stimulus. And axons were blocked if their maximum membrane potential at the putative node of excitation was ≥ –44 mV, or their respective inactivation (h) gate was ≤ 0.01 for 25 for at least 90 % of the simulation time (Fig. A.2). A designation of “blocked” also required that no action potentials were observed at the axon’s termini other than an initial onset response (e.g., see Fig. 4A). All simulations for assessing a conduction block were repeated to visually confirm each state (i.e., quiescent, active, or blocked).

Average spike counts in the nerve were calculated using a weighted average. For a given population of A*β* or A*δ* fibers, we summed the number of action potentials recorded for each of their five subpopulations. Then, we summed the total number of axon potentials across all subpopulations with normalized weightings (of ∼0.05:0.25:0.4:0.25:0.05) based on their respective fiber counts (Fig. 5A). This average action potential rate served as a proxy for the perceived intensity of electrocutaneous stimulation ^71^.

### Modeling tES of neurons

Subthreshold modulatory effects of current-regulated tES on the cortex were estimated using a population of spiking pyramidal neurons. Each neuron consisted of two compartments, including a spherical soma (diameter = 10 μm) and a cylindrical apical dendrite (length = 700 μm, diameter = 1.2 μm). The two-compartment model was reduced to an equivalent leaky integrate-and-fire (LIF) neuron model,

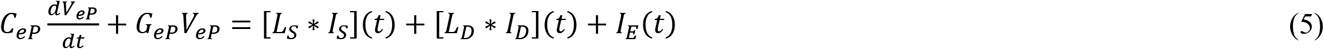

which we refer to as an extended point (eP) neuron to be consistent with its original derivation by Aspart et al. ^72^. *C*_*eP*_is the membrane capacitance, *G*_*eP*_is the membrane conductance, *L*_*S*_*\* I*_*S*_is a linear filter (*L*) convolved with the somatic (*S*) intracellular current (*I*_*S*_), *L*_*D*_*\* I*_*D*_is another linear filter convolved with the dendritic (*D*) intracellular current (*I*_*D*_), and *I*_*E*_is the equivalent intracellular current generated by an external applied electric field (E).

Linear filters were included in the eP neuron model to capture exactly the frequency-dependent input impedances of the soma and apical dendrite of the parent two-compartment neuron model. The linear filter for the soma is summarized by Equations 6 and 7:

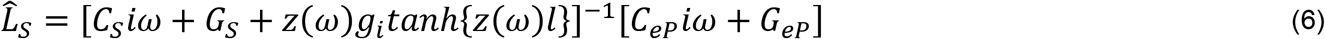

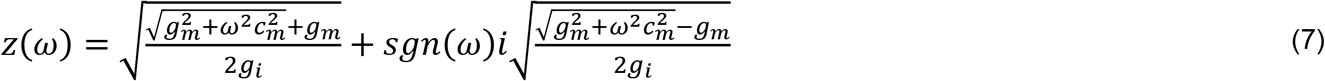

where, where *g*_*i*_is the intracellular conductance per unit length, *g*_*m*_is the membrane conductance per unit length, *c*_*m*_is the membrane capacitance per unit length, *l* is the length of the somal compartment, *ω* is angular frequency (2π*f*), *f* is frequency, *i* is the unit imaginary number, *sgn(*.) is the sign operator, and *^* denotes the Fourier transform. The linear filter for the apical dendrite is summarized by Equations 7 and 8:

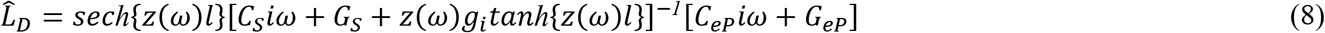

*C*_*eP*_ and *G*_*eP*_ were set equal to the somatic values in the parent model so that *C*_*eP*_ = *C*_*S*_ and *G*_*eP*_ = *G*_*S*_.

Synaptic noise (N), representing uncorrelated excitatory and inhibitory neural activity, was added to both the soma and apical dendrite using a Gauss-Markov process:

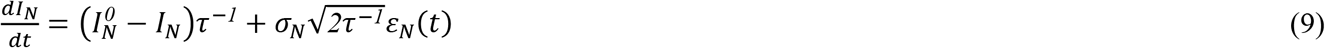

where, *I*_*N*_^*0*^ is the mean value of the noise, *σ*_N_ is the standard deviation of the noise, *τ* is the time constant of correlation, and *ε*_N_ is a standard Guassian process with zero mean and delta autocorrelation.

Finally, *I*_*E*_ was calculated based on the following transfer function:

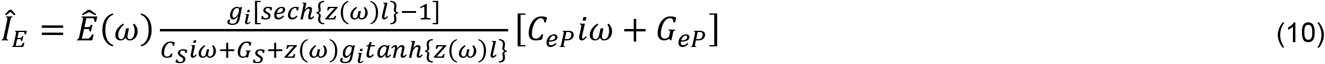

The magnitude of E was calculated using the seven-domain spherical head model described in the previous section. We assumed pyramidal neurons were located 1 mm below the cortical surface, and E was ideally colinear with the somato-dendritic axis of the eP neuron model. Equation 10 filters *E* by the input impedance of the pyramidal neuron (i.e., the second fractional factor in Equation 10) and then converts E to an equivalent intracellular current (*I*_*E*_) by further scaling by the eP model’s input admittance (i.e., the last factor in Equation 10). Details of the derivation of Equations 5–10 are summarized in another work ^72^. The model parameters used in the simulations are summarized in Table 1.

Simulations of pyramidal neuron firing were conducted in Python (version 3.7) using a Fourier approach. Applied currents and E were decomposed in the spectral domain using a fast Fourier Transform, multiplied by their respective linear filters (see Equations 6–8 and 10), and we used the inverse Fourier Transform to calculate the input time series for Equation 5. eP models were numerically solved using forward Euler time discretization. Uniform time steps of 100 μs were used in all cases except for stimulation frequencies greater than 1 kHz (or pulse widths less than 100 μs), where 1 μs was used. Simulations were conducted using 100 independent neurons over 10 s of simulated time.

We used the time-scale independent synchronization metric proposed by Kruez et al. ^75^ to analyze the synchronization of spike times in the pyramidal neuron population; the metric is bounded between 0 and 1, where 0 is complete desynchronization (no coincidence) of spike times, 1 is perfect synchronization (coincidence) of spike times, and coincidence is determined by comparing interspike intervals between neurons with interspike intervals within neurons. Population synchrony was analyzed for tACS, tPCS, and the combination of tDCS with tPCS. For tDCS, we also evaluated the average firing rate of the neuron population.

## Results

### *In Silico* Representation of tES

To study the nerve response to tES, we modeled 150 myelinated Aβ axons in a virtual cylindrical compartment within a semi-infinite medium (Fig. 1A). The position and size of the nerve were based on measurements from human supraorbital and supratrochlear nerves, and the conductivity and permittivity of the medium were frequency-dependent (Fig. 1B) ^76^. Electric fields generated in the volume conductor were combined with multi-compartment cable models of the axons to quantify nerve dynamics with different tES techniques, including transcranial direct current stimulation (tDCS), transcranial alternating current stimulation (tACS), transcranial pulsed current stimulation (tPCS), as well as combinations of these waveforms.

**Figure 1.**
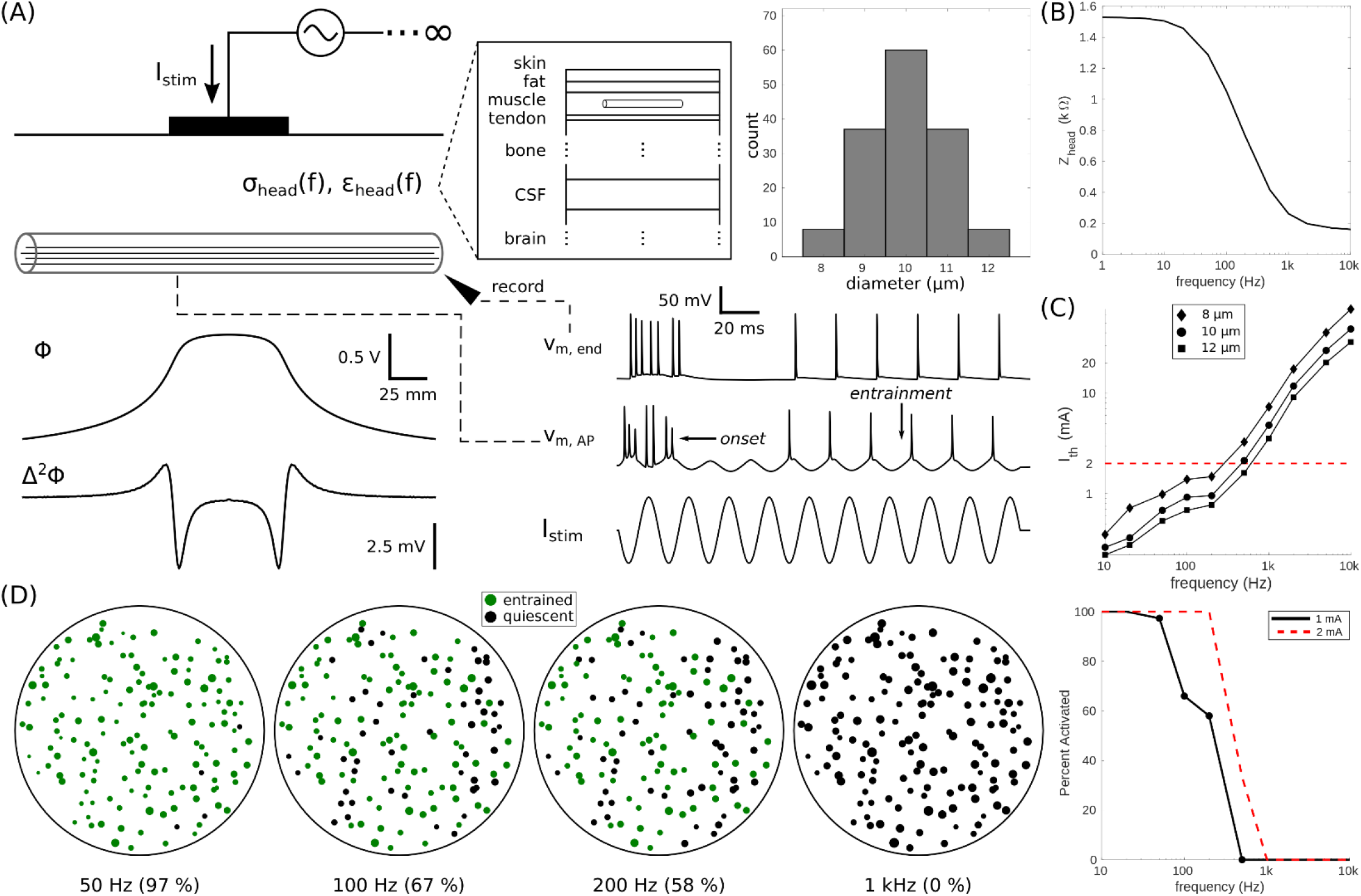
Axonal responses in the nerve are abolished at kilohertz frequencies. (A) The scalp model of transcranial electrical stimulation (tES) consisted of a disk electrode (diameter = 3 in) in contact with a semi-infinite “lossy” medium. The conductivity (σ) and permittivity (ε) of the medium reproduced the bulk head impedance of skin, soft tissues, and brain (see Methods and materials). The model nerve (diameter = 2 mm) was placed 6 mm below the surface of the electrode and consisted of 300 myelinated axons that were uniformly distributed. Fiber diameters of 8–12 μm were chosen to represent Aβ fibers. Sinusoidal currents were injected with the disk electrode to simulate transcranial alternating current stimulation (tACS). Axons that fired one action potential (AP) per phase of the stimulus were considered active / entrained. (B) Bode plot of head impedance (Z_head_) versus the frequency of the stimulus. (C) Strength-frequency curves of the stimulation current (I_stim_) required to activate a representative 10 μm axon at the nerve center. The standard limit of 2 mA for tACS is shown (*red dashed line*) for reference. (D) Percentage of axons that are entrained (*green*) or quiescent (*black*) during 1 mA tACS at four frequencies. Also shown are recruitment curves of percent axonal activation for all frequencies at 1 mA or 2 mA.

We also modeled endogenously firing pyramidal neurons to evaluate the immediate effects of tES on spike rate ^77^ and spike timing ^8,9^. Each neuron contained a soma and apical dendrite, into which picoamps of noisy current were injected to emulate endogenous synaptic activity (Fig. 2A), producing a basal firing rate of 2 Hz. The neuron model, although morphologically simplified, captured the general input-output properties of pyramidal neurons (Fig. 2B). The model pyramidal neuron was strongly filtered above hundreds of hertz ^78^, with a resonant frequency between 10 Hz and 20 Hz, and spike dynamics were modulated at electric fields of 1 V/m ^8,9^ (Fig. 2C/I). The combination of these two silico environments were used to analyze the temporal dynamics of active sham and sub-perception strategies for tES.

**Figure 2.**
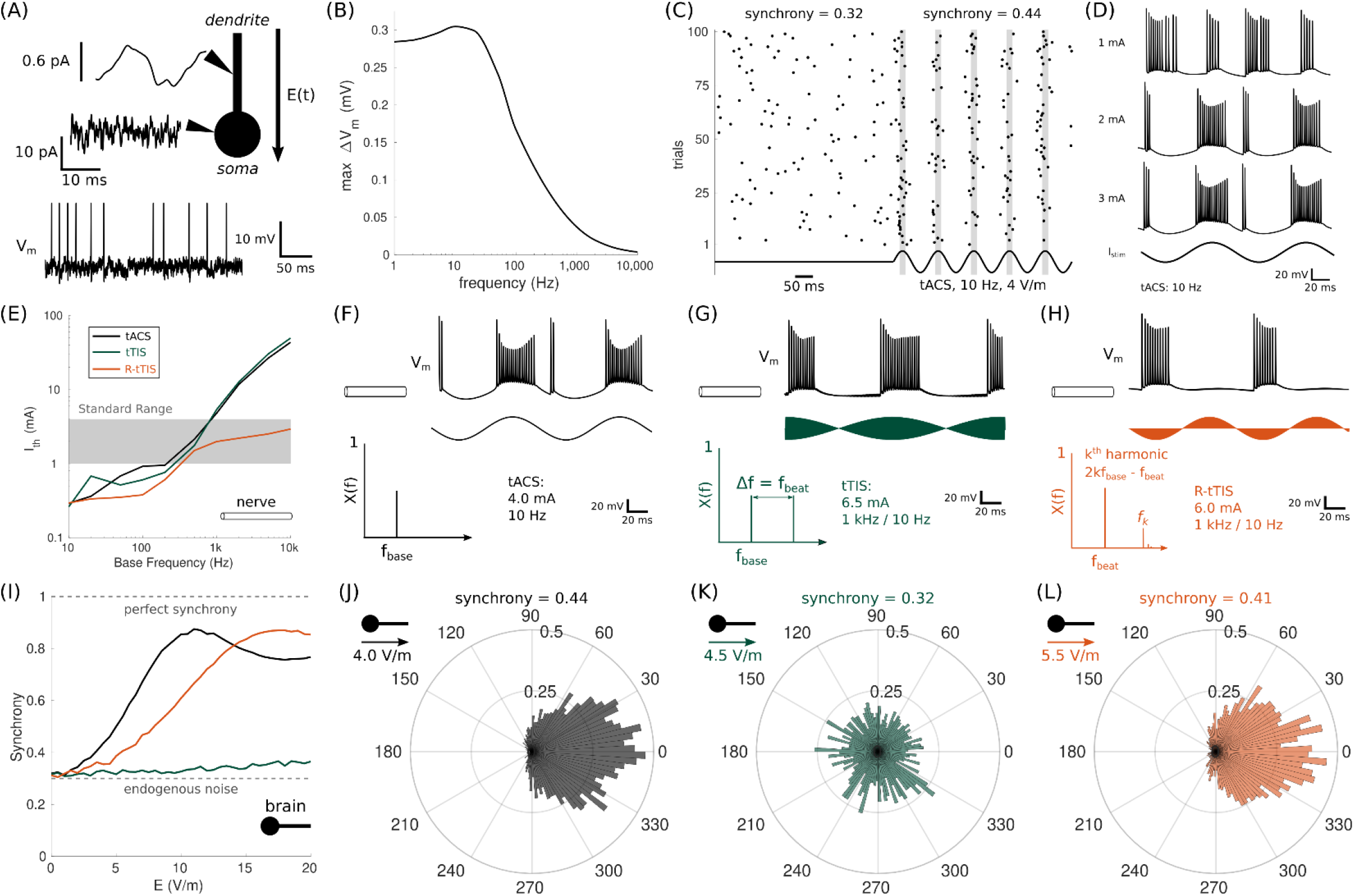
tTIS activates the nerve but not cortical tissue at kilohertz frequencies. (A) A model of a spiking pyramidal neuron with a spherical soma (diameter = 10 μm) and cylindrical apical dendrite (length = 700 μm, diameter = 1.2 μm). Synaptic activity in the neuron was simulated by injecting noisy intracellular currents. The intracellular currents followed a Gauss-Markov process with a mean of 6 pA and a standard deviation of 5 pA for the soma, and a mean of 0 pA and a standard deviation of 5 pA for the dendrite, altogether producing an average endogenous firing rate of approximately 2 Hz in the soma. (B) A Bode plot of the membrane response (ΔV_m_) without synaptic activity under an electric field of 1 V/m. (C) Spike synchronization (Synchrony) in a population of 100 pyramidal neurons without and with tACS. (D) Examples of axonal responses to tACS in the model nerve for increasing current amplitudes. (E) Strength-frequency curves of the threshold stimulation current (I_th_) required to activate a 10 μm axon located at the center of the model nerve for tACS, standard tTIS, and rectified tTIS (R-tTIS). The carrier and beat frequencies of tTIS and R-tTIS were 1 kHz and 10 Hz, respectively. The range of amplitudes explored in research studies (i.e., 1–4 mA) is highlighted in grey. (F–H) Examples of axonal membrane responses (V_m_) for tACS (*black*), tTIS (*green*), and R-tTIS (*orange*). The stimulation amplitudes were chosen to achieve approximate equivalence in the number of action potentials generated by each waveform. X(f) is the frequency spectrum of each respective waveform. (I) Synchrony of 100 pyramidal neurons for increasing electric field strengths. (I–L) Aggregate distribution of phase delays in the interspike intervals across 100 pyramidal neurons for the respective cases in F–H. 0° corresponds to an interspike interval of one over the stimulation frequency. The electric field magnitudes were calculated 1 mm below the brain surface in the tissue model (Fig. 1A).

### Filtering Properties of the Tissue Abate Axonal Responses

The first set of silico experiments were conducted with tACS. Previous reports demonstrate a marked reduction in cutaneous sensations and phosphenes above 100 Hz ^52,53^, despite axons being able to follow electrical stimuli up to several kilohertz ^79^. We postulated that this reduction in side effects occurs from filtering properties of the nervous tissue. The head impedance declined after 20 Hz and plateaued to a minimal value beyond 1 kHz (Fig. 1B). Myelinated axons faithfully responded to the tACS waveform (Fig. 2D) but required currents of greater than 2 mA for kilohertz frequencies (Fig. 1C). None of the myelinated axons were blocked at any frequencies below 10 mA. Therefore, this “dropout” effect in axonal responses (Fig. 1D) could be explained by a combination of passive and active filtering from the tissues and neural membranes, respectively. Impedance precipitously declined beyond 20 Hz, so less voltage was required to drive current through the medium, reducing the extracellular driving force for membrane polarization. Above 1 kHz, the head impedance plateaued, but activation thresholds still increased (Fig. 1B/C) due to only partial recovery (de-inactivation) of the sodium channels during persistent polarization.

The “dropout” effect observed in our nerve model motivated a second set of simulations to test if high frequency tACS could modulate spiking in pyramidal neurons (Fig. 2C) without concomitant activation of the cutaneous nerves. tACS at 10 Hz served as the base case, and we tested premixed tTIS waveforms: one, standard (amplitude-modulated) tTIS, and two, rectified tTIS (R-tTIS), which introduces a true low-frequency carrier at the beat frequency. The beat frequency for all AC waveforms was 10 Hz unless specified otherwise. Activation thresholds primarily increased with an increased carrier frequency but not the beat frequency (Fig. 2E), so larger currents were required for kilohertz tTIS and R-tTIS to match the nerve response of low-frequency tACS (Fig. 2F–H). Despite a larger dose, 1 kHz tTIS, did not appreciably synchronize pyramidal neurons at field strengths of ≤ 20 V/m because of the absence of any true low frequencies in its spectrum (Fig. 2I–K). However, R-tTIS had true low-frequency content at its beat frequency, so the modulatory strength of tACS and R-tTIS were comparable at matched beat frequencies (Fig. 2L). Yet, rectification of the kilohertz waveform did not completely eliminate its high-frequency carrier, so R-tTIS was less potent than tACS.

### Pulsatile and Direct Currents Limit Axonal Firing Rates

The next set of simulations focused on the principle that the intensity of electrocutaneous stimulation depends on the aggregate action potential count across all axons ^71^. tACS couldn’t modulate pyramidal neurons without concomitant activation of the cutaneous nerve, but we propose that the number of action potentials generated can be limited with pulsatile currents. First, we evaluated the comparative performance of tPCS and tACS at 10 Hz (Fig. 3A–C). tPCS was either monophasic with a positive phase, or biphasic with an additional negative phase, both of which were aligned with the extrema of tACS. Thresholds for axonal activation approached their rheobase at approximately 10 ms (Fig. 3D). Biphasic tPCS had lower activation thresholds for axons than monophasic tPCS due the additional cathodic phase (Fig. 3D), but the negative phase had no appreciable effect on modulating the pyramidal neurons for the waveforms tested (Fig. 3C). Synchronization of pyramidal neuron spiking with millisecond tPCS was possible below 10 mA, with shorter pulses requiring larger currents to achieve parity with tACS (Fig. 3E–H), and with matched modulatory effects, there was less variability in the spike timing compared to tACS (Fig3. I–K).

**Figure 3.**
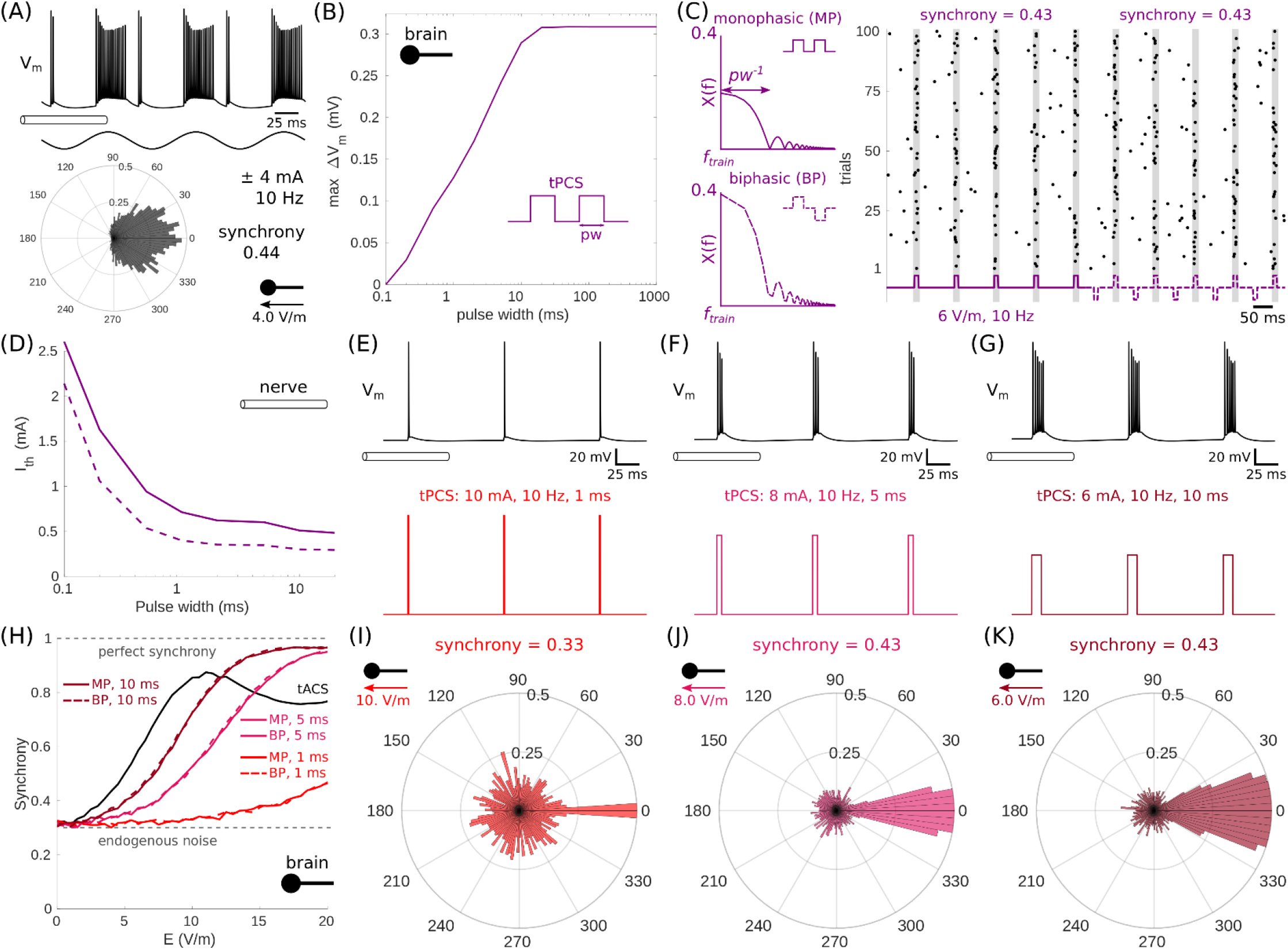
Pulsed currents synchronize action potentials stronger than tACS but at higher amplitudes. (A) Simulated action potentials in the nerve (*top*) and synchronization of spikes in 100 pyramidal neurons (*below*) for the base case, tACS at 10 Hz. (B) Strength-duration curve of the maximum somal membrane polarization (ΔV_m_) in the model pyramidal neuron for transcranial pulsed current stimulation (tPCS) with increasing duration. (C) Frequency spectra and raster plots of spike times for monophasic (MP, *solid line*) and biphasic (BP, *dashed line*) tPCS. The negative phase of the BP waveform occurred midway between cycles to match the timing and sign of tACS at a matched frequency. (D) Strength-frequency curves of the threshold stimulation current (I_th_) required to activate a 10 μm axon located at the center of the model nerve for MP and BP tPCS. (E–G) Examples of axonal membrane responses (V_m_) for tPCS at 1 ms (*red*), 5 ms (*pink*), and 10 ms (*maroon*). The stimulation amplitudes were chosen to achieve the same synchrony as the base case in A. However, current amplitudes were limited to 10 mA. (H) Synchrony of 100 pyramidal neurons for increasing electric field strengths with tPCS and tACS. (I–K) Aggregate distribution of phase delays in the interspike intervals across 100 pyramidal neurons for the respective cases in E–G. 0° corresponds to an interspike interval of one over the stimulation frequency. The electric field magnitudes were calculated 1 mm below the brain surface in the tissue model (Fig. 1A).

We also tested a strategy of combining pulsatile and direct currents to mitigate axonal firing, with the idea being that superposition of tPCS and tDCS would generate fewer action potentials than tPCS due to inactivation of sodium channels from sustained depolarization ^80^. tDCS generated three responses in the nerve: quiescence with or without a transient onset response, direct activation of the axon with periodic firing, and block from inactivation of sodium channels (Fig. 4A–B). The majority of large axons (Fig. 1A) were blocked at or above 3 mA (Fig. 4C), so we combined tPCS with 3 mA offset. Superposition of tPCS with 3 mA tDCS generated no additional action potentials in inactivated axons (Fig. 4D–F) while concomitantly synchronizing the pyramidal neurons at 10 mA or less (Fig. 4G–I).

**Figure 4.**
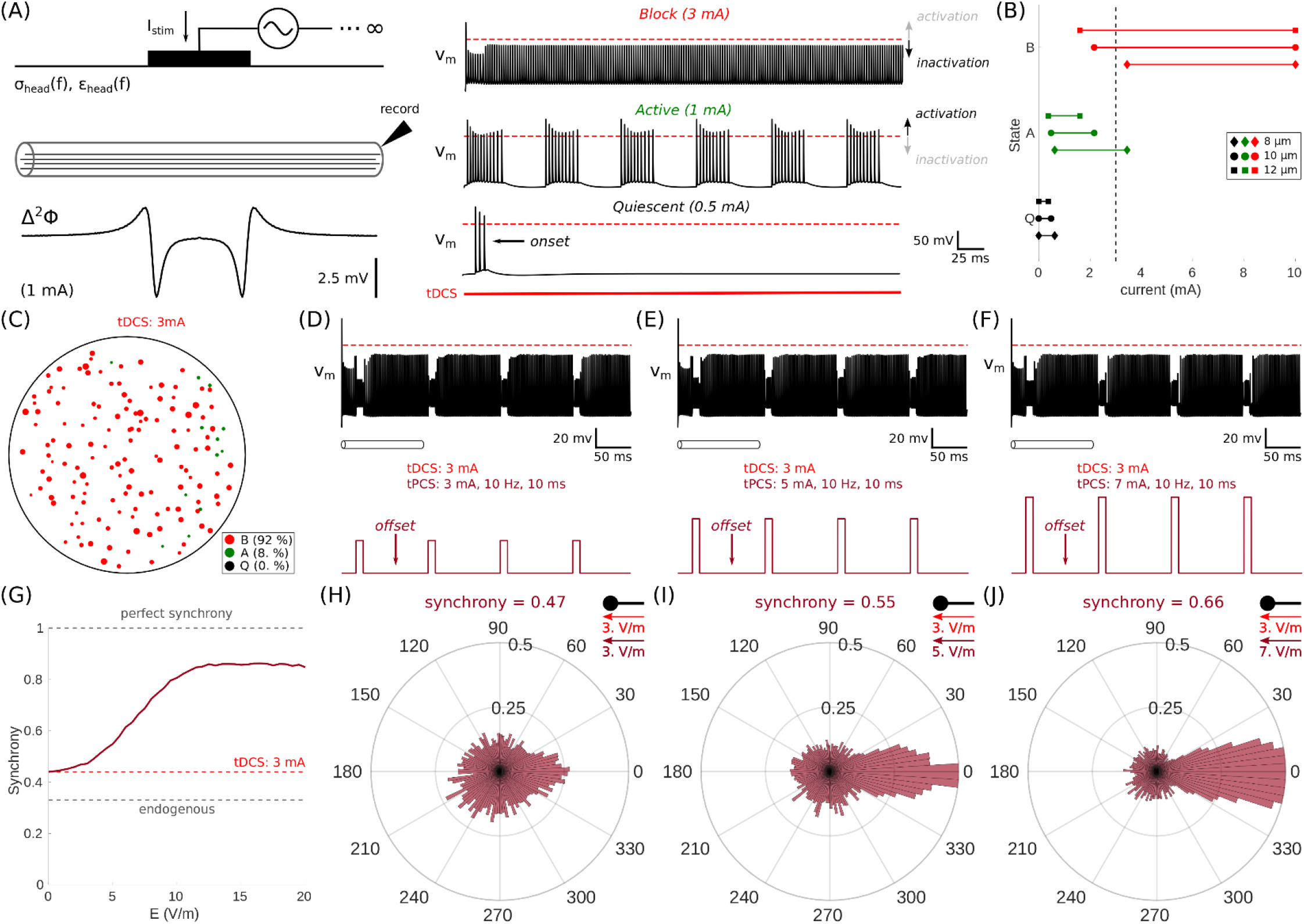
Conduction of action potentials is blocked at relatively high tDCS amplitudes. (A) The scalp model of tES (Fig. 1A) was used to evaluate nerve responses to tDCS. Axons exhibited three types of responses: quiescence (Q, *black*) with or without an onset response, persistent activation (A, *green*), and conduction block (B, *red*) with persistent sodium channel inactivation (see Appendix A). (B) State plots for three different fiber diameters at increasing current intensities. (C) Percentage of axons that are either Quiescent, Active, or Blocked for tDCS at 3 mA. (D–F) Examples of axonal membrane responses (V_m_) for a combination of tDCS at 3 mA, and tPCS at 3 mA (*left*), 5 mA (*middle*), or 7 mA (*right*). (H) Synchrony of 100 pyramidal neurons for increasing electric field strengths with 3 mA tDCS, or a combination of 3 mA tDCS and tPCS at increasing electric field strengths. (I–K) Aggregate distribution of phase delays in the interspike intervals across 100 pyramidal neurons for the respective cases in D–F. 0° corresponds to an interspike interval of one over the stimulation frequency. The electric field magnitudes were calculated 1 mm below the brain surface in the tissue model (Fig. 1A).

Cutaneous sensations reported during tES include tingling, itching, burning, and other painful sensations ^81^, which implicates activation of both large (8–12 μm) Aβ fibers and small (2–6 μm) Aδ fibers. Therefore, we modeled an additional population of small axons to quantify co-recruitment of these two populations in the nerve (Fig. 5A). As in prior simulations, tACS and tPCS were delivered at 10 Hz, and we used 10 ms as the representative case for tPCS. tACS had the overall highest rate of action potential generation across both populations (Fig. 5B). Below 2 mA, tDCS generated more action potentials than tPCS for both small and large axons, but above 2 mA, the onset of conduction block reduced the action potential rate more than the effect of limiting the stimulus duration with tPCS. These results provide a possible mechanistic explanation as to why tDCS is tolerated up to 4 mA compared to other techniques ^54^.

**Figure 5.**
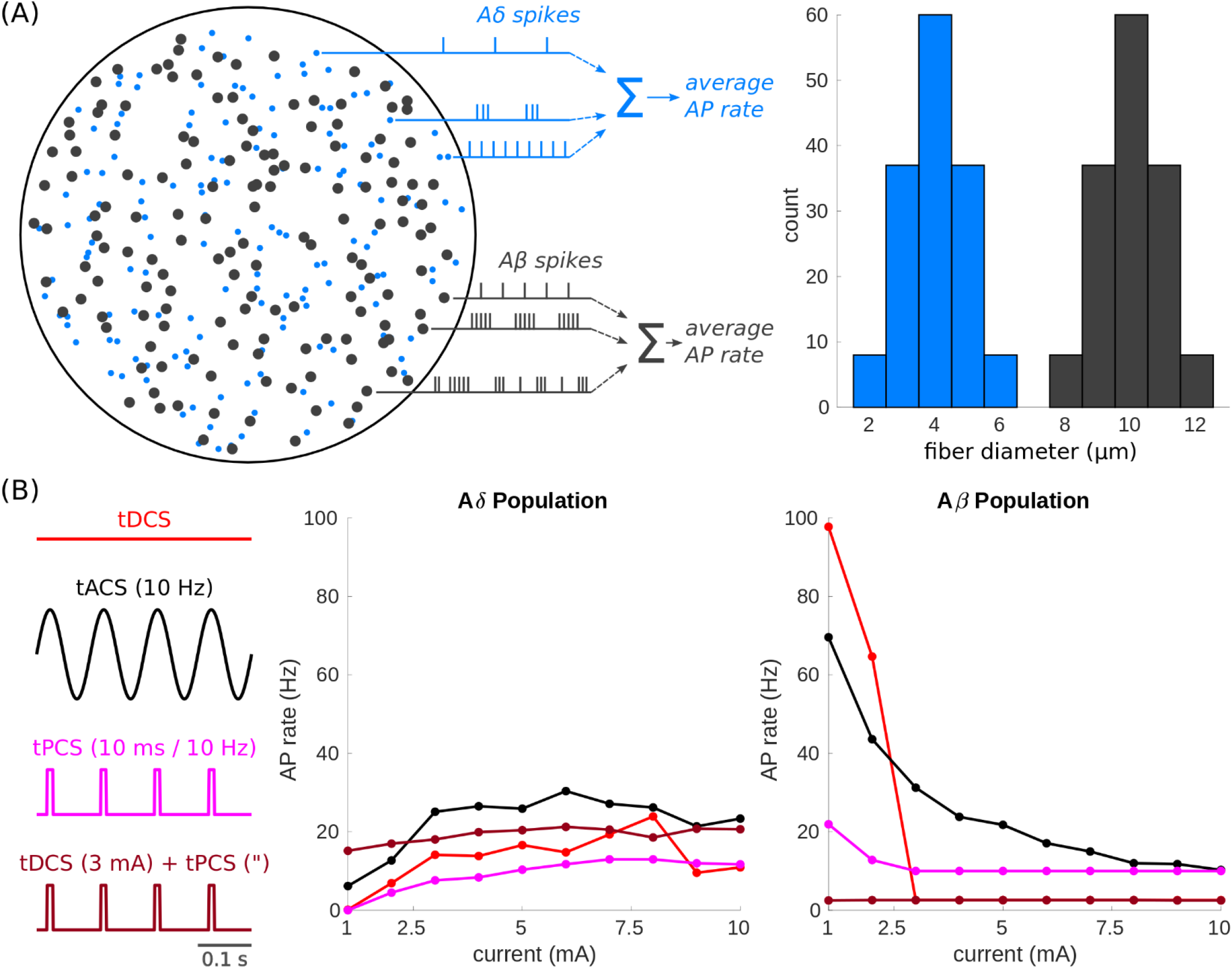
Recruitment dynamics in the nerve differ by fiber size and tES waveform. (A) The scalp model of tES (Fig. 1A) was used to quantify the average firing rates of action potentials (APs) across two different populations of 300 axons. Large axons with fiber diameters of 8–12 μm represented Aβ fibers (*black*). Small axons with fiber diameters of 2–6 μm represented Aδ fibers (*blue*). The number of APs was summed across each population separately and averaged by the respective number of axons. (B) The AP rates were calculated for tDCS (*red*), tACS at 10 Hz (*black*), 10 ms tPCS at 10 Hz (*pink*), and the combination of 3 mA tDCS and tPCS (*maroon*) at increasing current intensities. For the combination of tDCS and tPCS, current denotes the amplitude of tPCS with respect to the DC offset (e.g., see Fig. 4H).

### Kilohertz tACS Mimic Varied Cutaneous Responses with Minimal Cortical Modulation

Voltage-gated ion channels recover a portion of the low-frequency envelope from an amplitude-modulated wave through current rectification ^82^. Otherwise, the subthreshold low-pass filtering properties of the neural membrane predominate (Fig. 2B). This dichotomy in polarization dynamics at the scalp and cortical levels means that it’s theoretically possible to drive cutaneous nerve activation with negligible cortical modulation, permitting active sham stimulation for tACS (Fig. 2F,G, and I–K), and tDCS. tDCS below 2 mA primarily activated the nerve (Fig. 4B), with action potentials emerging in bursts between refractory periods to establish a stable limit cycle (Fig. 4A and 6A). At 1 mA tDCS, action potentials fired at approximately 14 Hz, and this response was emulated using 1 kHz tTIS at 6.8 mA with a matched beat frequency (Fig. 6A–B). At or above 3 mA tDCS, the majority of large axons were blocked (Fig. 4C), and we used 10 kHz tACS at 9.3 mA to match this response (Fig. 6C–D). In this case, 10 kHz tACS generated no action potentials because it’s threshold was significantly above 10 mA (Fig. 1C), and the amplitude was chosen to match the input electrical power, or ohmic heating, of 3 mA tDCS for parity. In all active sham settings, the cutaneous nerve model was activated with no appreciable modulation of the model pyramidal neurons (Fig. 6E–G).

**Figure 6.**
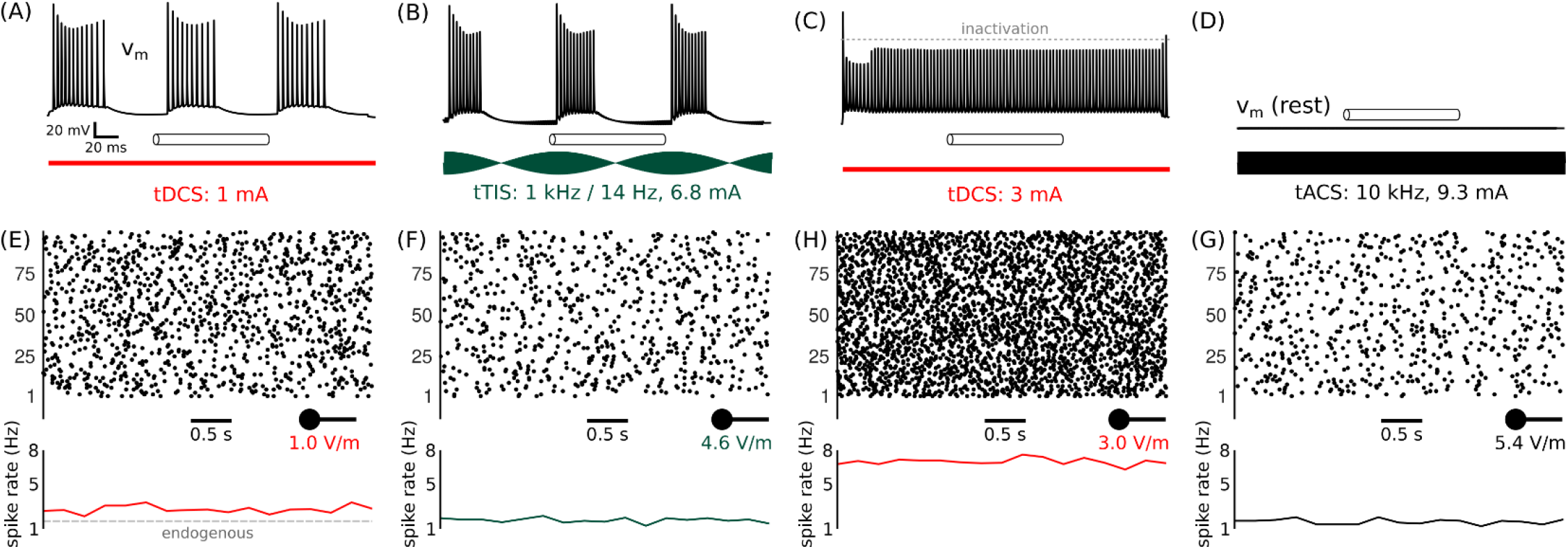
Kilohertz tACS emulates the nerve response of tDCS without cortical modulation. (A) Representative axonal response in the model nerve to anodal tDCS at 1 mA. The axon is at the nerve center with a fiber diameter of 10 μm. (B) The response in A was emulated using standard tTIS at 6.8 mA with a 1 kHz carrier frequency and a 14 Hz beat frequency. (C) Representative axonal response in the model nerve to anodal tDCS at 3 mA. The axon is quiescent due to the onset of the conduction block. (D) The nerve response in C was emulated using 10 kHz tACS at 9.3 mA. Axons were quiescent in both instances, so cases were matched based on the input electrical power (i.e., P = I^2^R). (E–H) Tive-averaged spike rate in 100 model pyramidal neurons for the respective cases in A–D. Spike counts averaged over 0.25 s bins. The endogenous firing rate is shown in E (*dashed grey line*). Electric fields were sampled 1 mm below the brain surface (see Fig. 1A).

## Discussion

This study dissected the biophysics underlying cutaneous responses to tES. Guided by the principle that the total spike count in afferent fibers dictates the perceived intensity of electrocutaneous stimulation ^71,83^, we identified three dynamical factors: 1) dielectric dispersion, 2) axonal refractoriness, and 3) conduction block, that could reconcile seemingly contradictory and non-monotonic sensory responses observed across different tES studies. Beyond hundreds of hertz, axonal responses to tACS were abolished first by the onset of alpha dispersions in the tissue medium, and at the kilohertz range, axonal refractoriness dominated, elevating excitation thresholds well above 10 mA. tDCS, on the other hand, blocked conduction in axons at currents above 2 mA, opening the possibility of combining tPCS or tACS with a DC block to achieve stronger synchronization of cortical spiking while mitigating cutaneous responses. Moreover, this study is the first to demonstrate the novel concept of using interferential tES as an active sham control. By leveraging differences in filtering effects at the scalp and brain levels, kilohertz tTIS could mimic the nerve responses of tDCS and tACS. This was accomplished by adjusting the amplitude and frequencies of the interferential wave, while simultaneously avoiding modulation of neural spiking in the cortex, even at field strengths above 10 V/m. All together, these insights provide a rigorous conceptual model for the design and analysis of new techniques for tES.

Our “dropout” hypothesis agrees with various experimental observations. The standard limit for tES is 2 mA for clinical studies, but this limit is likely overly conservative for certain waveforms. One study ^52^ looking at sub-kilohertz tACS reported that the severity of cutaneous responses at 250 Hz were no different than that of sham stimulation, and phosphenes were imperceivable at and above 140 Hz. Additionally, reports show that 5 kHz tACS is well-tolerated ^84^, with only transient, mild sensations above 5 mA ^53^. We posit that the observations of Turi et al. ^52^ are the product of alpha dispersions ^85^, which makes tissues more conductive at higher frequencies ^86,87^. Smaller head impedances drive more current with less voltage, so the field strength per unit current is diminished (Fig. 1D). Beyond 1 kHz, alpha dispersions plateau (Fig. 1B), and ion-channel dynamics are the limiting factor. Our original theory was that high-amplitude kilohertz tACS would block larger axons ^88^, but this was not observed at 10 mA or less (Fig. 1C). Rather, voltage-gated sodium channels in axons were only partially deactivated, elevating activation thresholds with increasing frequency.

Conduction block, however, was observed with direct current. tDCS at 4 mA has been well-tolerated in clinical studies with pain scores increasing only marginally from 1–4 mA ^54,89^, and in one cohort, the sensations of 4 mA tDCS and sham stimulation were equally severe ^55^. These observations implicate a non-monotonic recruitment trend (Fig. 4). Aβ fibers are initially activated up to 2 mA. However, beyond 2 mA, sodium channel inactivation in these large fibers leads to a conduction block ^80^, while smaller Aδ fibers are increasingly co-activated (Fig. 6). This is consistent with tingling being the most common sensation evoked by tDCS (in ∼75 % of individuals) from Aβ activation, following by itching, burning, and pricking (in ∼30 %) ^90^ from activation of nociceptive fibers ^91,92^. While it’s still unknown how sensory and nociceptive sensations are weighted in the overall unpleasantness of tDCS across individuals ^93^; tingling generally carries the largest weighting in self-reports of pain scores ^29^. Therefore, electrocutaneous inactivation of Aβ may help reduce discomfort during tDCS.

tPCS may be a suitable alternative to tDCS or tACS (Fig. 3A–C) depending on the application. Long-duration tPCS of hundred of milliseconds can emulate the modulatory effects of tDCS and with less adverse effects ^45,94,95^ (Fig. A.1), which agrees with our predictions below 2 mA (Fig. 6B). Yet, above 2 mA, conduction block is also a factor, and we posit that tDCS will be more tolerable than tPCS due to less overall nerve activation ^83^. Short-duration tPCS of 1–10 ms, on the other hand, is more suitable for emulating tACS (Fig. 3D–K). Because shorter stimulus durations decrease the perceived intensity of electrical stimulation for a set frequency ^71^, tPCS may be more tolerable than tACS, particularly when combined with a DC offset that blocks Aβ fibers (Fig. 4A– C) while simultaneously facilitating central sensory adaptation ^96^. Nonetheless, introduction of a DC offset may counteract the modulatory effects of tPCS at the network level when the constituents’ amplitudes are equally weighted ^50,97^, so the viability of this technique will depend on its tolerability for different amplitude weightings.

Transcranial temporal interfering stimulation, or tTIS ^46^, has also been proposed as an alternative to tACS. By generating regions of interference between two or more kilohertz current sources, tTIS may simultaneously block and activate neurons depending on the degree of interference ^47,82^. However, this requires direct activation of neurons, which is likely not possible at the human scale within tolerable stimulation limits ^37,98^. Spatial multiplexing with multiple independent current sources can theoretically boost the strength of tTIS ^73,99^, but even so, kilohertz tTIS still requires significant amplification to compensate for low-pass filtering through the neural membrane (Fig. 2B and 2I) ^100^. Although the modulatory effects of tTIS could arise from other biological mechanisms ^101–103^, the polarization strength of tTIS is expected to be less than that of tACS(Fig. 2) ^47,74^.

We propose a novel idea of using tTIS for active sham stimulation. Because the modulatory strength of tTIS depends primarily on the carrier frequency ^47^, its administration can be simplified by premixing the interferential waveform with the hardware, which we term premixed tTIS. This approach leverages the low-pass filtering properties of neurons to minimize brain modulation during concomitant nerve activation (Figs. 2 and 6), providing a means to explore the putative effects of nerve activation in isolation ^25,26^. Instead of minimizing skin responses with dermatological pretreatments ^28,29^, tTIS could be theoretically configured to mimic the nerve response of tACS (Fig. 2), tDCS (Fig. 6), or by extension, tPCS, by applying a window function to the stimulus. Better active placebo controls would help identify sources of variability across studies ^17^, and tTIS could be configured to study the effects of tES with minimal nerve activation (Fig. 1D), minimal cortical modulation (Fig. 2G), or both (Fig. 6D). Therefore, tTIS provides an invaluable tool to study the indirect rate-dependent effects of weak electric fields of 1–20 V/m on blood flow ^104^, the blood-brain barrier ^105^, or ion channels ^106^.

## Limitations

The models used in this study were spatially homogenized. Distributions of electric potentials vary across different individuals and brain regions due to variations in the material properties of the soft tissues and brain ^7,107^. Despite these simplifications, the models of the nerve and head were sized based on average measurements from adult humans, producing electric field magnitudes of at least 1 V/m with 1 mA of current applied across the scalp (Figs. 2J–L, 3I–K, 4H–J), consistent with electrical measurements in humans ^108^. This study focused on the temporal neural dynamics underlying the direct cutaneous response to tES, so we put more weight on studying the temporal dynamics of biophysical interactions between different tES waveforms and the filtering properties of neural tissues. While the exact applied currents required to evoke neural responses will vary across individuals and brain regions, the geometric simplifications made herein are not strictly prohibitive to studying the biophysical principles underlying cutaneous responses to tES.

We also simplified the neuroanatomy of our model cortical neurons. Biophysics dictates that a large nearby neuron with axes collinear with the electric field’s orientation will be the most responsive to extracellular electrical stimuli, so our analyses focused on large, superficial pyramidal neurons. Our models recapitulate general input-output properties of pyramidal neurons in the cortex ^72^, and can be modulated at electric field intensities of ∼1 V/m and above (Figs. 2J–L, 3I–K, 4H–J), consistent with *in vivo* recording in macaques ^8,109^. However, we did not model the varied types, sizes, orientations, and morphologies of neural elements that reside in cortical tissues ^110^. tES of heterogeneous cortical tissue produces differential modulatory effects at the mesoscopic (multi-neuron) and macroscopic (network) levels ^111^, and the general tenets of anodic tES increasing neural activity, and cathodic tES decreasing neural activity are less reliable over large spatial and temporal scales. Nonetheless, these principles at the single-neuron level are still useful for comparing the direct modulatory effects of varied tES waveforms for first-principles analysis.

## Conclusions

Our results support a new conceptual model for tES based on established biophysical principles. The tolerability of tES is likely not fixed, but rather, varies across different waveforms. DC currents are likely the most tolerable due, in part, to the onset of conduction block with increasing current amplitudes, and these blocking stimuli can be combined with pulsatile currents to achieve potentially higher doses with an overall less severe cutaneous response. Moreover, we propose a novel application of kilohertz tTIS, where premixed interferential waveforms are used to activate nearby nerves while fully leveraging the low-pass filtering properties of neurons to avoid concomitant modulation of cortical neurons.

## Acknowledgements

This work was supported by grants from the National Institute of Neurological Disorders and Stroke (R01NS105690) and the National Institute of Mental Health (R01MH102238) of the National Institutes of Health. This work made use of the High-Performance Computing Resource in the Core Facility for Advanced Research Computing at Case Western Reserve University.

## Supporting Information

Figure S1 shows that pulsatile currents can mimic the modulatory effects of direct currents. Figure S2 shows that the membrane potential and h-gate values predict the onset of a conduction block.

## References

1. Brittain J-S, Probert-Smith P, Aziz TZ, Brown P. Tremor suppression by rhythmic transcranial current stimulation. Current Biology. 2013;23(5):436–440.

2. Del Felice A, Castiglia L, Formaggio E, et al. Personalized transcranial alternating current stimulation (tACS) and physical therapy to treat motor and cognitive symptoms in Parkinson’s disease: A randomized cross-over trial. NeuroImage: Clinical. 2019;22:101768. doi:10.1016/j.nicl.2019.101768

3. Ganguly J, Murgai AA, Sharma S, Aur D, Jog MS. Non-invasive transcranial electrical stimulation in movement disorders. Frontiers in neuroscience. 2020;14:522.

4. Brunoni AR, Moffa AH, Fregni F, et al. Transcranial direct current stimulation for acute major depressive episodes: Meta-analysis of individual patient data. British Journal of Psychiatry. 2016;208(6):522–531. doi:10.1192/bjp.bp.115.164715

5. Razza LB, Palumbo P, Moffa AH, et al. A systematic review and meta-analysis on the effects of transcranial direct current stimulation in depressive episodes. Depression and Anxiety. 2020;37(7):594–608. doi:10.1002/da.23004

6. Shivakumar V, Dinakaran D, Narayanaswamy JC, Venkatasubramanian G. Noninvasive brain stimulation in obsessive–compulsive disorder. Indian journal of psychiatry. 2019;61(Suppl 1):S66.

7. Huang Y, Liu AA, Lafon B, et al. Measurements and models of electric fields in the in vivo human brain during transcranial electric stimulation. Ivry R, ed. eLife. 2017;6:e18834. doi:10.7554/eLife.18834

8. Johnson L, Alekseichuk I, Krieg J, et al. Dose-dependent effects of transcranial alternating current stimulation on spike timing in awake nonhuman primates. Sci Adv. 2020;6(36):eaaz2747. doi:10.1126/sciadv.aaz2747

9. Krause MR, Vieira PG, Csorba BA, Pilly PK, Pack CC. Transcranial alternating current stimulation entrains single-neuron activity in the primate brain. Proc Natl Acad Sci USA. 2019;116(12):5747. doi:10.1073/pnas.1815958116

10. Bikson M, Grossman P, Thomas C, et al. Safety of Transcranial Direct Current Stimulation: Evidence Based Update 2016. Brain Stimul. 2016;9(5):641–661. doi:10.1016/j.brs.2016.06.004

11. Cucca A, Sharma K, Agarwal S, Feigin AS, Biagioni MC. Tele-monitored tDCS rehabilitation: feasibility, challenges and future perspectives in Parkinson’s disease. Journal of NeuroEngineering and Rehabilitation. 2019;16(1):20. doi:10.1186/s12984-019-0481-4

12. Hordacre B. The Role of Telehealth to Assist In-Home tDCS: Opportunities, Promising Results and Acceptability. Brain Sciences. 2018;8(6). doi:10.3390/brainsci8060102

13. Riggs A, Patel V, Paneri B, Portenoy RK, Bikson M, Knotkova H. At-home transcranial direct current stimulation (tDCS) with telehealth support for symptom control in chronically-ill patients with multiple symptoms. Frontiers in behavioral neuroscience. 2018;12:93.

14. Horvath JC, Carter O, Forte JD. Transcranial direct current stimulation: five important issues we aren’t discussing (but probably should be). Front Syst Neurosci. 2014;8:2–2. doi:10.3389/fnsys.2014.00002

15. Krause B, Cohen Kadosh R. Not all brains are created equal: the relevance of individual differences in responsiveness to transcranial electrical stimulation. Frontiers in Systems Neuroscience. 2014;8:25. doi:10.3389/fnsys.2014.00025

16. López-Alonso V, Cheeran B, Río-Rodríguez D, Fernández-del-Olmo M. Inter-individual Variability in Response to Non-invasive Brain Stimulation Paradigms. Brain Stimulation: Basic, Translational, and Clinical Research in Neuromodulation. 2014;7(3):372–380. doi:10.1016/j.brs.2014.02.004

17. Fonteneau C, Mondino M, Arns M, et al. Sham tDCS: A hidden source of variability? Reflections for further blinded, controlled trials. Brain Stimulation. 2019;12(3):668–673. doi:10.1016/j.brs.2018.12.977

18. Ambrus GG, Al-Moyed H, Chaieb L, Sarp L, Antal A, Paulus W. The fade-in – Short stimulation – Fade out approach to sham tDCS – Reliable at 1 mA for naïve and experienced subjects, but not investigators. Brain Stimulation: Basic, Translational, and Clinical Research in Neuromodulation. 2012;5(4):499–504. doi:10.1016/j.brs.2011.12.001

19. Miranda PC, Faria P, Hallett M. What does the ratio of injected current to electrode area tell us about current density in the brain during tDCS? Clinical Neurophysiology. 2009;120(6):1183–1187. doi:10.1016/j.clinph.2009.03.023

20. Gandiga PC, Hummel FC, Cohen LG. Transcranial DC stimulation (tDCS): A tool for double-blind sham-controlled clinical studies in brain stimulation. Clinical Neurophysiology. 2006;117(4):845–850. doi:10.1016/j.clinph.2005.12.003

21. De Smet S, Nikolin S, Moffa A, et al. Determinants of sham response in tDCS depression trials: a systematic review and meta-analysis. Progress in Neuro-Psychopharmacology and Biological Psychiatry. 2021;109:110261. doi:10.1016/j.pnpbp.2021.110261

22. Greinacher R, Buhôt L, Möller L, Learmonth G. The time course of ineffective sham-blinding during low‐intensity (1 mA) transcranial direct current stimulation. European Journal of Neuroscience. 2019;50(8):3380–3388.

23. Russo R, Wallace D, Fitzgerald PB, Cooper NR. Perception of Comfort During Active and Sham Transcranial Direct Current Stimulation: A Double Blind Study. Brain Stimulation. 2013;6(6):946–951. doi:10.1016/j.brs.2013.05.009

24. Ezquerro F, Moffa AH, Bikson M, et al. The Influence of Skin Redness on Blinding in Transcranial Direct Current Stimulation Studies: A Crossover Trial. Neuromodulation: Technology at the Neural Interface. 2017;20(3):248–255. doi:10.1111/ner.12527

25. Asamoah B, Khatoun A, Mc Laughlin M. tACS motor system effects can be caused by transcutaneous stimulation of peripheral nerves. Nature Communications. 2019;10(1):266. doi:10.1038/s41467-018-08183-w

26. Liu A, Vöröslakos M, Kronberg G, et al. Immediate neurophysiological effects of transcranial electrical stimulation. Nature Communications. 2018;9(1):5092. doi:10.1038/s41467-018-07233-7

27. Rizzo V, Terranova C, Crupi D, Sant’angelo A, Girlanda P, Quartarone A. Increased Transcranial Direct Current Stimulation After Effects During Concurrent Peripheral Electrical Nerve Stimulation. Brain Stimulation: Basic, Translational, and Clinical Research in Neuromodulation. 2014;7(1):113–121. doi:10.1016/j.brs.2013.10.002

28. Guarienti F, Caumo W, Shiozawa P, et al. Reducing Transcranial Direct Current Stimulation-Induced Erythema With Skin Pretreatment: Considerations for Sham-Controlled Clinical Trials. Neuromodulation: Technology at the Neural Interface. 2015;18(4):261–265. doi:10.1111/ner.12230

29. McFadden JL, Borckardt JJ, George MS, Beam W. Reducing procedural pain and discomfort associated with transcranial direct current stimulation. Brain Stimulation. 2011;4(1):38–42. doi:10.1016/j.brs.2010.05.002

30. Fischer DB, Fried PJ, Ruffini G, et al. Multifocal tDCS targeting the resting state motor network increases cortical excitability beyond traditional tDCS targeting unilateral motor cortex. NeuroImage. 2017;157:34–44. doi:10.1016/j.neuroimage.2017.05.060

31. Neri F, Mencarelli L, Menardi A, et al. A novel tDCS sham approach based on model-driven controlled shunting. Brain Stimulation. 2020;13(2):507–516. doi:10.1016/j.brs.2019.11.004

32. Richardson JD, Fillmore P, Datta A, Truong D, Bikson M, Fridriksson J. Toward development of sham protocols for high-definition transcranial direct current stimulation (HD-tDCS). NeuroRegulation. 2014;1(1):62–62.

33. Ruffini G, Fox MD, Ripolles O, Miranda PC, Pascual-Leone A. Optimization of multifocal transcranial current stimulation for weighted cortical pattern targeting from realistic modeling of electric fields. NeuroImage. 2014;89:216–225. doi:10.1016/j.neuroimage.2013.12.002

34. Chew T, Ho K-A, Loo CK. Inter- and Intra-individual Variability in Response to Transcranial Direct Current Stimulation (tDCS) at Varying Current Intensities. Brain Stimulation. 2015;8(6):1130–1137. doi:10.1016/j.brs.2015.07.031

35. Wiethoff S, Hamada M, Rothwell JC. Variability in Response to Transcranial Direct Current Stimulation of the Motor Cortex. Brain Stimulation. 2014;7(3):468–475. doi:10.1016/j.brs.2014.02.003

36. Laakso I, Tanaka S, Koyama S, De Santis V, Hirata A. Inter-subject Variability in Electric Fields of Motor Cortical tDCS. Brain Stimulation. 2015;8(5):906–913. doi:10.1016/j.brs.2015.05.002

37. Vöröslakos M, Takeuchi Y, Brinyiczki K, et al. Direct effects of transcranial electric stimulation on brain circuits in rats and humans. Nature Communications. 2018;9(1):483. doi:10.1038/s41467-018-02928-3

38. Colella M, Paffi A, De Santis V, Apollonio F, Liberti M. Effect of skin conductivity on the electric field induced by transcranial stimulation techniques in different head models. Physics in Medicine & Biology. 2021;66(3):035010.

39. Khadka N, Bikson M. Role of skin tissue layers and ultra-structure in transcutaneous electrical stimulation including tDCS. Physics in Medicine & Biology. 2020;65(22):225018.

40. Evans C, Bachmann C, Lee JSA, Gregoriou E, Ward N, Bestmann S. Dose-controlled tDCS reduces electric field intensity variability at a cortical target site. Brain Stimulation. 2020;13(1):125–136. doi:10.1016/j.brs.2019.10.004

41. Rich TL, Gillick BT. Electrode Placement in Transcranial Direct Current Stimulation—How Reliable Is the Determination of C3/C4? Brain Sciences. 2019;9(3). doi:10.3390/brainsci9030069

42. Woods AJ, Bryant V, Sacchetti D, Gervits F, Hamilton R. Effects of Electrode Drift in Transcranial Direct Current Stimulation. Brain Stimulation. 2015;8(3):515–519. doi:10.1016/j.brs.2014.12.007

43. Caulfield KA, Badran BW, DeVries WH, et al. Transcranial electrical stimulation motor threshold can estimate individualized tDCS dosage from reverse-calculation electric-field modeling. Brain Stimulation. 2020;13(4):961–969. doi:10.1016/j.brs.2020.04.007

44. Chaieb L, Paulus W, Antal A. Evaluating Aftereffects of Short-Duration Transcranial Random Noise Stimulation on Cortical Excitability. Chen R, ed. Neural Plasticity. 2011;2011:105927. doi:10.1155/2011/105927

45. Jaberzadeh S, Bastani A, Zoghi M. Anodal transcranial pulsed current stimulation: A novel technique to enhance corticospinal excitability. Clinical Neurophysiology. 2014;125(2):344–351. doi:10.1016/j.clinph.2013.08.025

46. Grossman N, Bono D, Dedic N, et al. Noninvasive Deep Brain Stimulation via Temporally Interfering Electric Fields. Cell. 2017;169(6):1029-1041.e16. doi:10.1016/j.cell.2017.05.024

47. Howell B, McIntyre CC. Feasibility of Interferential and Pulsed Transcranial Electrical Stimulation for Neuromodulation at the Human Scale. Neuromodulation: Technology at the Neural Interface. 2020;n/a(n/a). doi:10.1111/ner.13137

48. Neudorfer C, Chow CT, Boutet A, et al. Kilohertz-frequency stimulation of the nervous system: A review of underlying mechanisms. Brain Stimulation: Basic, Translational, and Clinical Research in Neuromodulation. 2021;14(3):513–530. doi:10.1016/j.brs.2021.03.008

49. Murphy OW, Hoy KE, Wong D, Bailey NW, Fitzgerald PB, Segrave RA. Transcranial random noise stimulation is more effective than transcranial direct current stimulation for enhancing working memory in healthy individuals: Behavioural and electrophysiological evidence. Brain Stimulation: Basic, Translational, and Clinical Research in Neuromodulation. 2020;13(5):1370–1380. doi:10.1016/j.brs.2020.07.001

50. Thibaut A, Russo C, Morales-Quezada L, et al. Neural signature of tDCS, tPCS and their combination: Comparing the effects on neural plasticity. Neuroscience Letters. 2017;637:207–214. doi:10.1016/j.neulet.2016.10.026

51. Ambrus GG, Paulus W, Antal A. Cutaneous perception thresholds of electrical stimulation methods: Comparison of tDCS and tRNS. Clinical Neurophysiology. 2010;121(11):1908–1914. doi:10.1016/j.clinph.2010.04.020

52. Turi Z, Ambrus GG, Janacsek K, et al. Both the cutaneous sensation and phosphene perception are modulated in a frequency-specific manner during transcranial alternating current stimulation. Restor Neurol Neurosci. 2013;31(3):275–285. doi:10.3233/RNN-120297

53. Kunz P, Antal A, Hewitt M, Neef A, Opitz A, Paulus W. 5 kHz Transcranial Alternating Current Stimulation: Lack of Cortical Excitability Changes When Grouped in a Theta Burst Pattern. Frontiers in Human Neuroscience. 2017;10:683. doi:10.3389/fnhum.2016.00683

54. Khadka N, Borges H, Paneri B, et al. Adaptive current tDCS up to 4 mA. Brain Stimulation. 2020;13(1):69–79. doi:10.1016/j.brs.2019.07.027

55. Workman CD, Kamholz J, Rudroff T. The Tolerability and Efficacy of 4 mA Transcranial Direct Current Stimulation on Leg Muscle Fatigability. Brain Sciences. 2020;10(1). doi:10.3390/brainsci10010012

56. Greenberg RJ, Velte TJ, Humayun MS, Scarlatis GN, De Juan E. A computational model of electrical stimulation of the retinal ganglion cell. IEEE Transactions on Biomedical Engineering. 1999;46(5):505–514.

57. Wang B, Petrossians A, Weiland JD. Reduction of edge effect on disk electrodes by optimized current waveform. IEEE Transactions on Biomedical Engineering. 2014;61(8):2254–2263.

58. Wiley JD, Webster JG. Analysis and control of the current distribution under circular dispersive electrodes. IEEE Transactions on Biomedical Engineering. 1982;(5):381–385.

59. Chopra K, Calva D, Sosin M, et al. A comprehensive examination of topographic thickness of skin in the human face. Aesthetic surgery journal. 2015;35(8):1007–1013.

60. Choi Y-J, Lee K-W, Gil Y-C, Hu K-S, Kim H-J. Ultrasonographic analyses of the forehead region for injectable treatments. Ultrasound in medicine & biology. 2019;45(10):2641–2648.

61. Farzana F, Shah BA, Shahdad S, ul Haq PZ, Sarmast A, Ali Z. Computed tomographic scanning measurement of skull bone thickness: a single center study. International Journal of Research in Medical Sciences. 2018;6(3):913.

62. Vatta F, Meneghini F, Esposito F, Mininel S, Di Salle F. Realistic and spherical head modeling for EEG forward problem solution: a comparative cortex-based analysis. Comput Intell Neurosci. 2010;2010:972060–972060. doi:10.1155/2010/972060

63. Gabriel S, Lau R, Gabriel C. The dielectric properties of biological tissues: III. Parametric models for the dielectric spectrum of tissues. Physics in medicine and biology. 1996;41(11):2271.

64. Baumann SB, Wozny DR, Kelly SK, Meno FM. The electrical conductivity of human cerebrospinal fluid at body temperature. IEEE Transactions on Biomedical Engineering. 1997;44(3):220–223. doi:10.1109/10.554770

65. Butson CR, McIntyre CC. Tissue and electrode capacitance reduce neural activation volumes during deep brain stimulation. Clin Neurophysiol. 2005;116(10):2490–2500. doi:10.1016/j.clinph.2005.06.023

66. Hines ML, Carnevale NT. Neuron: A Tool for Neuroscientists. The Neuroscientist. 2001;7(2):123–135. doi:10.1177/107385840100700207

67. McIntyre CC, Richardson AG, Grill WM. Modeling the excitability of mammalian nerve fibers: influence of afterpotentials on the recovery cycle. Journal of Neurophysiology. 2002;87(2):995–1006.

68. Howell B, McIntyre CC. Analyzing the tradeoff between electrical complexity and accuracy in patientspecific computational models of deep brain stimulation. J Neural Eng. 2016;13(3):036023. doi:10.1088/1741-2560/13/3/036023

69. Jacobs JM. On internodal length. Journal of Anatomy. 1988;157:153–162.

70. Rydmark M, Berthold C-H. Electron microscopic serial section analysis of nodes of Ranvier in lumbar spinal roots of the cat: a morphometric study of nodal compartments in fibres of different sizes. Journal of neurocytology. 1983;12(4):537–565.

71. Graczyk EL, Schiefer MA, Saal HP, Delhaye BP, Bensmaia SJ, Tyler DJ. The neural basis of perceived intensity in natural and artificial touch. Science Translational Medicine. 2016;8(362):362ra142. doi:10.1126/scitranslmed.aaf5187

72. Aspart F, Ladenbauer J, Obermayer K. Extending Integrate-and-Fire Model Neurons to Account for the Effects of Weak Electric Fields and Input Filtering Mediated by the Dendrite. PLOS Computational Biology. 2016;12(11):e1005206. doi:10.1371/journal.pcbi.1005206

73. Rampersad S, Roig-Solvas B, Yarossi M, et al. Prospects for transcranial temporal interference stimulation in humans: A computational study. NeuroImage. 2019;202:116124. doi:10.1016/j.neuroimage.2019.116124

74. Negahbani E, Kasten FH, Herrmann CS, Fröhlich F. Targeting alpha-band oscillations in a cortical model with amplitude-modulated high-frequency transcranial electric stimulation. NeuroImage. 2018;173:3–12. doi:10.1016/j.neuroimage.2018.02.005

75. Kreuz T, Chicharro D, Houghton C, Andrzejak RG, Mormann F. Monitoring spike train synchrony. Journal of neurophysiology. 2013;109(5):1457–1472.

76. Gabriel S, Lau RW, Gabriel C. The dielectric properties of biological tissues: II. Measurements in the frequency range 10 Hz to 20 GHz. Physics in Medicine and Biology. 1996;41(11):2251.

77. Kronberg G, Rahman A, Sharma M, Bikson M, Parra LC. Direct current stimulation boosts hebbian plasticity in vitro. Brain Stimulation. 2020;13(2):287–301. doi:10.1016/j.brs.2019.10.014

78. Vaidya SP, Johnston D. Temporal synchrony and gamma-to-theta power conversion in the dendrites of CA1 pyramidal neurons. Nature Neuroscience. 2013;16(12):1812–1820. doi:10.1038/nn.3562

79. Crosby ND, Janik JJ, Grill WM. Modulation of activity and conduction in single dorsal column axons by kilohertz-frequency spinal cord stimulation. Journal of Neurophysiology. 2016;117(1):136–147. doi:10.1152/jn.00701.2016

80. Bhadra N, Kilgore KL. Direct current electrical conduction block of peripheral nerve. IEEE Transactions on Neural Systems and Rehabilitation Engineering. 2004;12(3):313–324.

81. Kessler SK, Turkeltaub PE, Benson JG, Hamilton RH. Differences in the experience of active and sham transcranial direct current stimulation. Brain Stimul. 2012;5(2):155–162. doi:10.1016/j.brs.2011.02.007

82. Mirzakhalili E, Barra B, Capogrosso M, Lempka SF. Biophysics of Temporal Interference Stimulation. Cell Systems. 2020;11(6):557-572.e5.

83. Martinsen ØG, Grimnes S, Piltan H. Cutaneous perception of electrical direct current. ITBM-RBM. 2004;25(4):240–243. doi:10.1016/j.rbmret.2004.09.012

84. Chaieb L, Antal A, Pisoni A, et al. Safety of 5 kHz tACS. Brain Stimulation: Basic, Translational, and Clinical Research in Neuromodulation. 2014;7(1):92–96. doi:10.1016/j.brs.2013.08.004

85. H. P. Schwan. Electrical properties of tissues and cell suspensions: mechanisms and models. In: Proceedings of 16th Annual International Conference of the IEEE Engineering in Medicine and Biology Society. Vol 1.; 1994:A70–A71 vol.1. doi:10.1109/IEMBS.1994.412155

86. Cole KS, Cole RH. Dispersion and Absorption in Dielectrics I. Alternating Current Characteristics. The Journal of Chemical Physics. 1941;9(4):341–351. doi:10.1063/1.1750906

87. Cole KS, Cole RH. Dispersion and Absorption in Dielectrics II. Direct Current Characteristics. The Journal of Chemical Physics. 1942;10(2):98–105. doi:10.1063/1.1723677

88. Bhadra N, Lahowetz EA, Foldes ST, Kilgore KL. Simulation of high-frequency sinusoidal electrical block of mammalian myelinated axons. Journal of Computational Neuroscience. 2007;22(3):313–326. doi:10.1007/s10827-006-0015-5

89. Chhatbar PY, Chen R, Deardorff R, et al. Safety and tolerability of transcranial direct current stimulation to stroke patients–A phase I current escalation study. Brain stimulation. 2017;10(3):553–559.

90. Poreisz C, Boros K, Antal A, Paulus W. Safety aspects of transcranial direct current stimulation concerning healthy subjects and patients. Brain Research Bulletin. 2007;72(4):208–214. doi:10.1016/j.brainresbull.2007.01.004

91. Beissner F, Brandau A, Henke C, et al. Quick Discrimination of Adelta and C Fiber Mediated Pain Based on Three Verbal Descriptors. PLOS ONE. 2010;5(9):e12944. doi:10.1371/journal.pone.0012944

92. Ringkamp M, Schepers RJ, Shimada SG, et al. A role for nociceptive, myelinated nerve fibers in itch sensation. Journal of Neuroscience. 2011;31(42):14841–14849.

93. Fertonani A, Ferrari C, Miniussi C. What do you feel if I apply transcranial electric stimulation? Safety, sensations and secondary induced effects. Clinical Neurophysiology. 2015;126(11):2181–2188. doi:10.1016/j.clinph.2015.03.015

94. Groppa S, Bergmann TO, Siems C, Mölle M, Marshall L, Siebner HR. Slow-oscillatory transcranial direct current stimulation can induce bidirectional shifts in motor cortical excitability in awake humans. Neuroscience. 2010;166(4):1219–1225. doi:10.1016/j.neuroscience.2010.01.019

95. Jaberzadeh S, Bastani A, Zoghi M, Morgan P, Fitzgerald PB. Anodal Transcranial Pulsed Current Stimulation: The Effects of Pulse Duration on Corticospinal Excitability. PLOS ONE. 2015;10(7):e0131779. doi:10.1371/journal.pone.0131779

96. Graczyk EL, Delhaye BP, Schiefer MA, Bensmaia SJ, Tyler DJ. Sensory adaptation to electrical stimulation of the somatosensory nerves. Journal of Neural Engineering. 2018;15(4):046002. doi:10.1088/1741-2552/aab790

97. Thibaut A, Russo C, Hurtado-Puerto AM, et al. Effects of Transcranial Direct Current Stimulation, Transcranial Pulsed Current Stimulation, and Their Combination on Brain Oscillations in Patients with Chronic Visceral Pain: A Pilot Crossover Randomized Controlled Study. Frontiers in Neurology. 2017;8:576. doi:10.3389/fneur.2017.00576

98. Alekseichuk I, Mantell K, Shirinpour S, Opitz A. Comparative modeling of transcranial magnetic and electric stimulation in mouse, monkey, and human. NeuroImage. 2019;194:136–148. doi:10.1016/j.neuroimage.2019.03.044

99. J. Cao, P. Grover. STIMULUS: Noninvasive Dynamic Patterns of Neurostimulation Using Spatio-Temporal Interference. IEEE Transactions on Biomedical Engineering. Published online 2019:1-1. doi:10.1109/TBME.2019.2919912

100. Esmaeilpour Z, Kronberg G, Reato D, Parra LC, Bikson M. Temporal interference stimulation targets deep brain regions by modulating neural oscillations. Brain Stimulation. 2021;14(1):55–65. doi:10.1016/j.brs.2020.11.007

101. Gellner A-K, Reis J, Fritsch B. Glia: A Neglected Player in Non-invasive Direct Current Brain Stimulation. Frontiers in Cellular Neuroscience. 2016;10:188. doi:10.3389/fncel.2016.00188

102. Wachter D, Wrede A, Schulz-Schaeffer W, et al. Transcranial direct current stimulation induces polarity-specific changes of cortical blood perfusion in the rat. Experimental Neurology. 2011;227(2):322–327. doi:10.1016/j.expneurol.2010.12.005

103. Zannou AL, Khadka N, FallahRad M, Truong DQ, Kopell BH, Bikson M. Tissue Temperature Increases by a 10 kHz Spinal Cord Stimulation System: Phantom and Bioheat Model. Neuromodulation: Technology at the Neural Interface. 2019;0(0). doi:10.1111/ner.12980

104. Turner DA, Degan S, Galeffi F, Schmidt S, Peterchev AV. Rapid, Dose-Dependent Enhancement of Cerebral Blood Flow by transcranial AC Stimulation in Mouse. Brain Stimulation. 2021;14(1):80–87. doi:10.1016/j.brs.2020.11.012

105. Shin DW, Fan J, Luu E, et al. In Vivo Modulation of the Blood–Brain Barrier Permeability by Transcranial Direct Current Stimulation (tDCS). Annals of Biomedical Engineering. 2020;48(4):1256–1270. doi:10.1007/s10439-020-02447-7

106. Pelletier SJ, Cicchetti F. Cellular and Molecular Mechanisms of Action of Transcranial Direct Current Stimulation: Evidence from In Vitro and In Vivo Models. International Journal of Neuropsychopharmacology. 2015;18(pyu047). doi:10.1093/ijnp/pyu047

107. Kasten FH, Duecker K, Maack MC, Meiser A, Herrmann CS. Integrating electric field modeling and neuroimaging to explain inter-individual variability of tACS effects. Nature Communications. 2019;10(1):5427. doi:10.1038/s41467-019-13417-6

108. Vöröslakos M, Takeuchi Y, Brinyiczki K, et al. Direct effects of transcranial electric stimulation on brain circuits in rats and humans. Nature Communications. 2018;9(1):483. doi:10.1038/s41467-018-02928-3

109. Krause MR, Vieira PG, Csorba BA, Pilly PK, Pack CC. Transcranial alternating current stimulation entrains single-neuron activity in the primate brain. Proc Natl Acad Sci USA. 2019;116(12):5747. doi:10.1073/pnas.1815958116

110. Traub RD, Contreras D, Cunningham MO, et al. Single-column thalamocortical network model exhibiting gamma oscillations, sleep spindles, and epileptogenic bursts. Journal of neurophysiology. 2005;93(4):2194–2232.

111. Tanaka T, Isomura Y, Kobayashi K, Hanakawa T, Tanaka S, Honda M. Electrophysiological Effects of Transcranial Direct Current Stimulation on Neural Activity in the Rat Motor Cortex. Frontiers in Neuroscience. 2020;14.

