## Supplemental Data for "Mimicking and mitigating the cutaneous response to transcranial electrical stimulation using interferential and combinatorial techniques"

**Supporting Information**

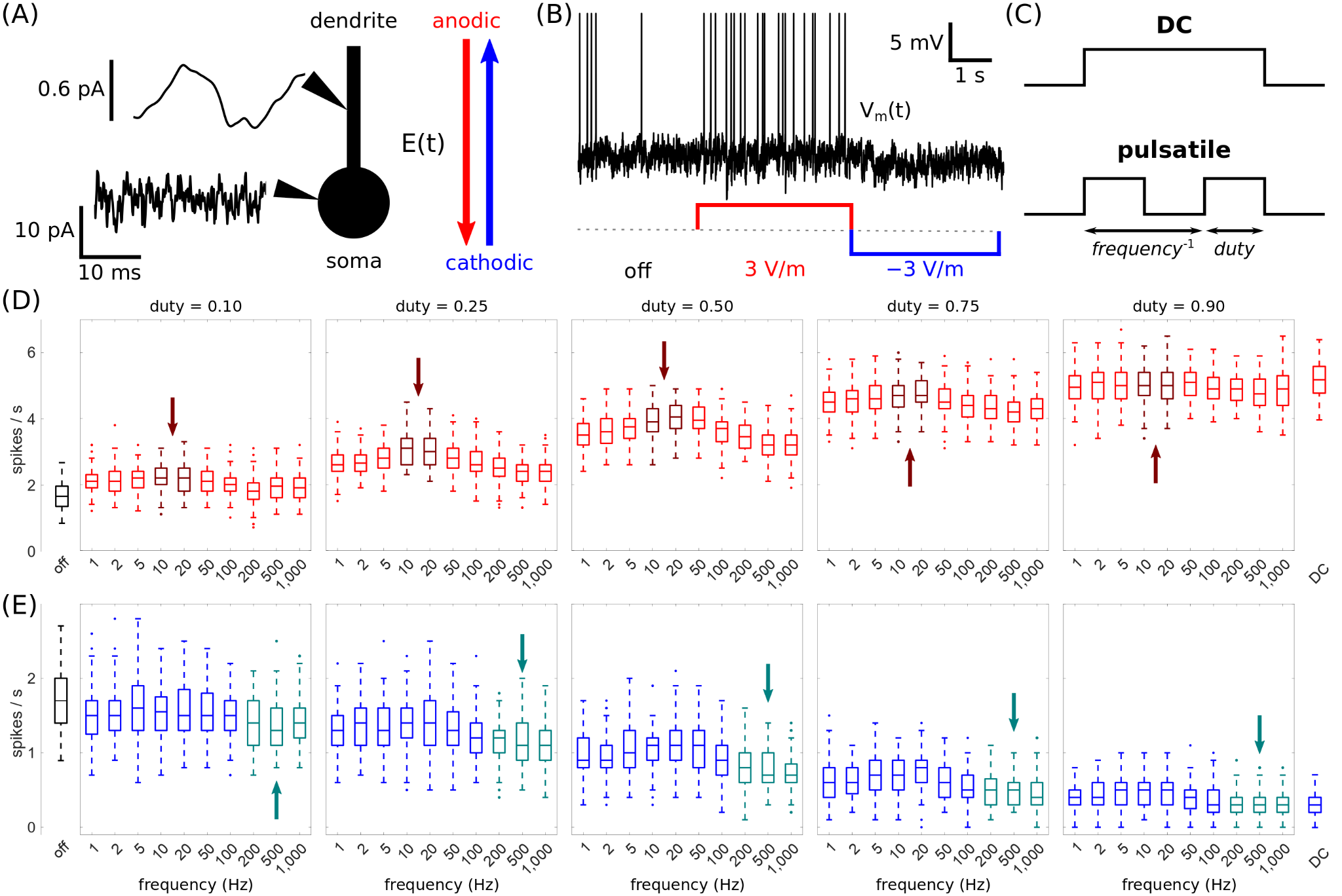

**Figure S1. Pulsatile currents can mimic the modulatory effects of direct currents.** (A) A model of a spiking pyramidal neuron with a spherical soma (diameter = 10 μm) and cylindrical apical dendrite (length = 700 μm, diameter = 1.2 μm). Synaptic activity in the neuron was simulated by injecting noisy intracellular currents. The intracellular currents followed a Gauss-Markov process with a mean of 6 pA and a standard deviation of 5 pA for the soma, and a mean of 0 pA and a standard deviation of 5 pA for the dendrite, altogether producing an average endogenous firing rate of approximately 2 Hz in the soma. Currents entering (exiting) the spherical cortex were defined as anodic (cathodic). (B) The model pyramidal neuron’s membrane potential (V_m_) with no stimulus, anodic tDCS, and cathodic tDCS. (C) Pulsatile currents are compared to direct current (DC) based on their frequency and duty cycle. (D) Average spike of the model pyramidal neuron over ten seconds for anodic pulsatile currents of increasing frequency and duty cycle. The spike rate is also shown for DC (*right*). The dark maroon boxes and arrows highlight a local peak in firing rates between 10 Hz and 20 Hz for each duty cycle. (E) The same as D except for cathodic pulsatile currents. The teal boxes and arrows highlight a local minimum in the firing rate at approximately 500 Hz for each duty cycle.

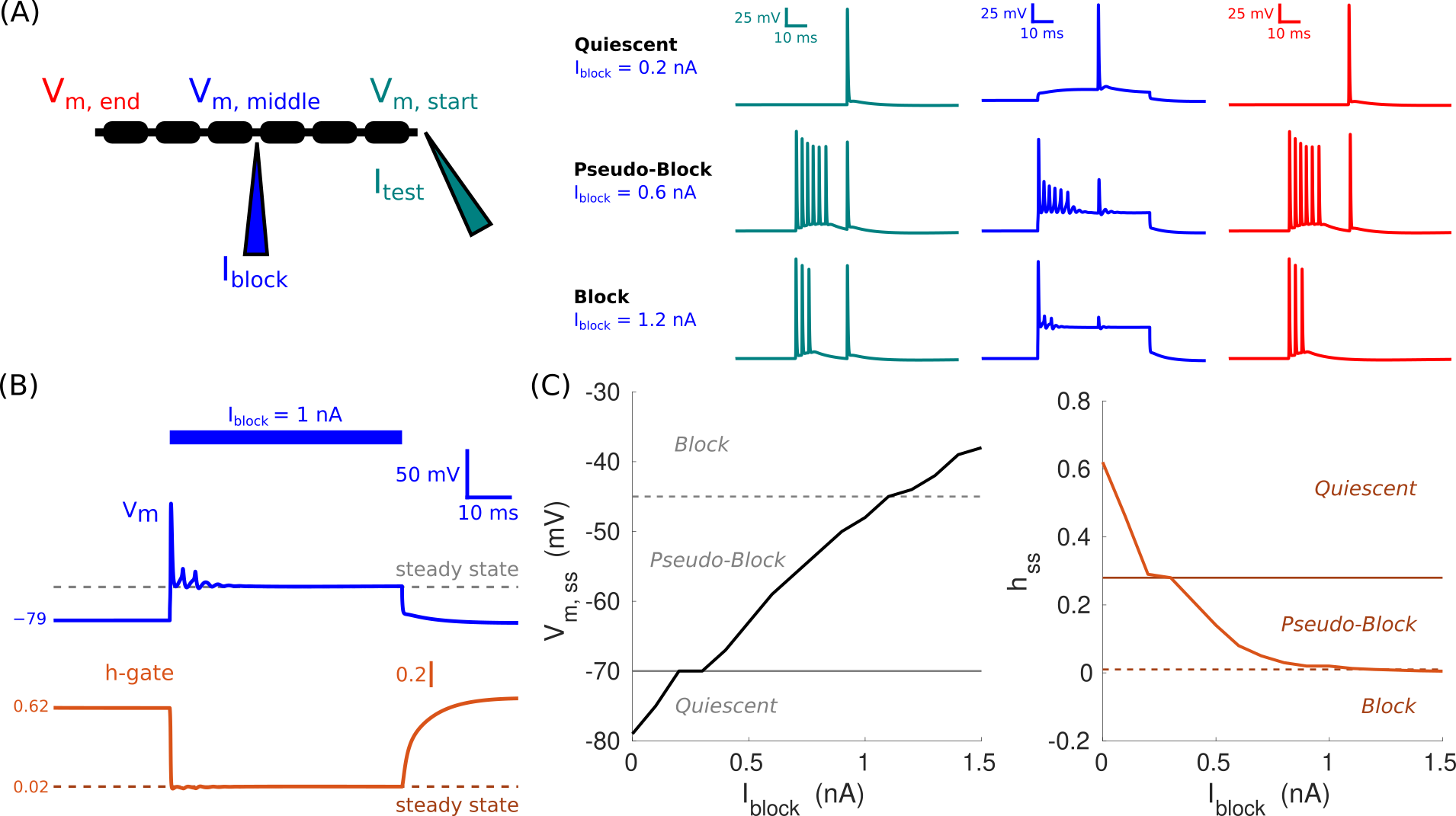

**Figure S2. The membrane potential and h-gate values predict the onset of a conduction block.**

(A) A model of a myelinated axon (fiber diameter = 10 μm, length = 120 mm) was stimulated using two intracellular currents. A block current (I_block_, *blue*) of variable amplitude was administered at the middle node (node 61) beginning at 25 ms for a duration of 50 ms. A test current (I_test_, *teal*) of 100 μs in duration and 1 nA in amplitude was administered at the first node to generate an action potential. The membrane potential (V_m_) was recorded at the start (*teal*), middle (*blue*), and end (*red*) of the axon to assess if a conduction block was present. Minus the first set of action potentials (i.e., onset response) generated by I_block_, the axon was considered pseudo-blocked if the test action potential arrived at the final node and blocked if not.

(B) V_m_ and the respective inactivation (h) gate parameter at the middle node for I_block_ of 1 nA. Steady-state values for V_m_ and h (*dashed lines*) were the respective average values over 50–75 ms.

(C) Steady-state (ss) values of V_m_ and h for increasing I_block_. There was an inverse relationship between V_m, ss_ and h_ss_. For V_m,ss_ (h_ss_), axons were quiescent below (above) the solid line, pseudo-blocked between the solid and dashed lines, and blocked above (below) the dashed line.
